# Ultrasensitive graphene FET aptasensor for direct attomolar detection of glutamate in human clinical samples

**DOI:** 10.1101/2025.11.05.686731

**Authors:** Mafalda Abrantes, Yolanda Blanco, Rafael P. Giacomazzi, Isabel P. Moreira, Patricia Monteiro, Jérôme Borme, Maria Vieira-Coelho, Sérgio F. Sousa, Carlos Briones, Luis Jacinto, Pedro Alpuim

**Affiliations:** International Iberian Nanotechnology Laboratory, 4715-330, Braga, Portugal; Physics Center of Minho and Porto Universities (CF-UM-UP), 4710-057, Braga, Portugal; Department of Biomedicine – Experimental Biology Unit, Faculty of Medicine, University of Porto (FMUP), 4200-319 Porto, Portugal; Department of Molecular Evolution, Centro de Astrobiología (CAB), CSIC-INTA, 28850 Torrejón de Ardoz, Madrid, Spain; LAQV/REQUIMTE, BioSIM—Department of Biomedicine, Faculty of Medicine, University of Porto (FMUP), 4200-319 Porto, Portugal; Neurology Department, Hospital Trofa Saúde, 4404-501 Vila Nova de Gaia, Portugal; Doctoral Program in Neurosciences, Faculty of Medicine, University of Porto (FMUP), 4200-319 Porto, Portugal; Rise-Health - Department of Biomedicine, Faculty of Medicine, University of Porto (FMUP), 4200-319 Porto, Portugal; Psychiatry Department, São João Local Health Unit (CHUSJ), 4200-319, Porto, Portugal; Department of Biomedicine – Pharmacology and Therapeutics Unit, Faculty of Medicine, University of Porto (FMUP), 4200-319 Porto, Portugal; Departament of Chemical Engineering, Faculty of Sciences and Engineering, University of Coimbra, 3030-790 Coimbra, Portugal

**Author notes:** These authors share senior authorship (Luis Jacinto, Pedro Alpuim). Corresponding author. Department of Molecular Evolution, Centro de Astrobiología (CAB), CSIC-INTA, 28850 Torrejón de Ardoz, Madrid, Spain. *Email address* (C. Briones). Corresponding author. Department of Biomedicine – Experimental Biology Unit, Faculty of Medicine, University of Porto (FMUP), 4200-319 Porto, Portugal. *Email address* (L. Jacinto). Corresponding author. International Iberian Nanotechnology Laboratory, 4715-330, Braga, Portugal. *Email address* (P. Alpuim).

## Abstract

Glutamate, the principal excitatory neurotransmitter in the brain, is crucial for cognition and memory, and its dysregulation is implicated in several neurological disorders, including Alzheimer’s Disease and epilepsy. However, precise and high-throughput quantification of glutamate in physiological samples remains challenging. Here, we report an ultrasensitive and highly specific glutamate aptamer-based biosensor, or aptasensor, developed through in silico design and microfabrication, followed by in vitro and clinical validation. The biosensor consists of arrays of graphene field-effect transistors functionalized with a novel DNA aptamer, NG-Apt-Glu, designed and characterized computationally and biochemically, revealing two putative glutamate binding sites. The aptasensor detects glutamate in artificial cerebrospinal fluid with a 1 aM detection limit, a wide linear range (1 aM–10 pM), and 24 mV/decade sensitivity, showing strong selectivity against GABA, glutamine, dopamine, and serotonin. To evaluate clinical applicability, glutamate levels were measured in cerebrospinal fluid from patients with Alzheimer’s Disease, showing a significant increase relative to controls and correlating with neurofilament light chain concentrations, a biomarker of neuronal death. These findings underscore glutamate’s involvement in Alzheimer’s pathophysiology and its potential as a biomarker for neurodegeneration. This ultrasensitive graphene-based aptasensor enables point-of-care monitoring, paving the way for early diagnosis and the development of novel therapeutic strategies.

## 1. Introduction

Neurological disorders such as Alzheimer’s Disease (AD) are increasingly prevalent [1], but lack effective molecular biomarkers that can contribute to early diagnosis or accelerate the discovery of novel therapies. One factor contributing to the onset and progression of many of these disorders is dysfunction of neuronal communication mediated by neurotransmitters [2]. Glutamate is the major excitatory neurotransmitter in the brain and is involved in several fundamental cerebral functions, including cognition and memory [3], [4]. Dysfunction of glutamate levels and dynamics has been correlated with excitotoxic effects implicated in the pathogenesis of AD and epilepsy [4], [5], or with excitatory/inhibitory imbalances, which may underlie brain disorders such as autism and schizophrenia [6].

Measuring glutamate levels in biological samples with high sensitivity, accuracy, and replicability in a high-throughput format is a significant challenge. Analytical methods available for clinical investigation are typically limited to enzyme-linked immunosorbent assays (ELISA) [7], [8], or high-precision liquid chromatography (HPLC) coupled with spectrofluorometric or mass spectrometric (MS) detection [9], [10]. Although these approaches have high selectivity, and the latter also shows high sensitivity, they can be very time-consuming and costly and have a long turnaround time for results. Hence, recent efforts have focused on developing small-scale biosensors, but balancing size, speed, sensitivity, and precision has proven difficult. Because glutamate is not electroactive at physiological pH [11], voltametric methods involving direct oxidation-reduction of the target molecule, widely used in biosensors for the detection of neurotransmitters such as dopamine and serotonin [12], cannot be applied. Thus, most glutamate biosensors have focused on indirect methods, mainly relying on enzymatic reactions that can create electroactive byproducts through the glutamate oxidase (GluOx) or the glutamate dehydrogenase (GLDH) enzymes [13], [14], [15]. Some enzyme-free methods have also been proposed, using the periplasmic glutamate-binding protein (GluBP) [16], metabotropic glutamate receptors (mGluR) [17], polymer synthetic receptors [18], [19], or catalytic nanoparticles [20], [21], [22], [23]. However, regardless of the biorecognition approach, the majority of these biosensors use voltammetric and amperometric methods for transduction, which may have disadvantages related to selectivity, low signal-to-noise ratio, and device complexity. Moreover, they typically suffer from low sensitivity and a short working range.

Field-effect transistor (FET)-based biosensors can enable ultrasensitive detection, even in complex samples, due to their ability to rapidly transduce small detection events [24], [25], but have been rarely applied to glutamate detection. The few previously reported examples, which included glutamate biorecognition via mGluR [17] or GluOx [26], [27], also showed limited sensitivity. This was due to the large-sized biorecognition elements used, which, upon detection, produced minor charge alterations that mostly fell outside the Debye length, the distance over which the electrolyte can screen charge alterations in liquid-gate FET sensors [28].

One solution to overcome the Debye length limitation in FET biosensors is the use of small-sized biorecognition elements, such as aptamers, which are typically short oligonucleotide sequences selected in vitro, whose 3D structure allows them to bind to the desired target molecules with high affinity and specificity [29], [30], [31], [32]. Because they are significantly smaller than enzymes and antibodies, when used as molecular probes for sensor functionalization, they can drive charge re-orientation upon target binding to occur within the Debye length [33]. However, selecting and biochemically characterizing novel aptamers for small molecular targets, such as neurotransmitters, is especially challenging [34]. In vitro selection of aptamers is typically performed using an iterative process called systematic evolution of ligands by exponential enrichment (SELEX), where a very large number of random RNA or ssDNA sequences are subjected to amplification/selection cycles to improve their binding affinity to the target molecule [35], [36]. This directed evolution process has been mainly used for large molecular weight targets such as proteins, which contain multiple potential aptamer-binding sites and show a significant mass differential between the bound complex and unbound aptamer [37]. To operate within the Debye length and retain sensors’ ultrasensitive performance, aptamers need to be as short as possible, which limits their conformational repertoire and the putative number of target-binding sites.

Recent modifications of the SELEX process, such as Capture SELEX [38] and electrochemical SELEX [39], have tried to address this limitation. However, although they are more cost-efficient than traditional SELEX, they have not yet been applied to selecting aptamers for neurotransmitter targets. An alternative, emerging approach to streamline rational aptamer development and reduce production cost and time is based on in silico optimization and computational predictions, with subsequent biochemical characterization of the most promising candidates [40]. In silico approaches can be used to both predict novel aptamer sequences by simulating the SELEX process through machine learning [41], as well as to rank and characterize aptamer candidates and aptamer-target binding sites through computational workflows that include 3D structure prediction, molecular and docking dynamics simulations, and binding energy calculations [40], [42], [43]. While the former is more challenging due to the lack of dedicated universal databases, the latter has been successfully used for optimal selection of previously identified aptamer candidates for clinically relevant targets [44], [45], [46].

Graphene aptasensors, which combine aptamers and graphene FETs (gFETs), have been previously described for ultrasensitive neurotransmitter sensing due to their unique electronic properties, stability in physiological environments, and high surface-to-area ratio for transduction [24], [47]. However, they have not yet been reported for glutamate sensing. Thus, in the present work, we combined in silico design and characterization with biochemical validation to select a novel glutamate aptamer, which, when coupled with highly sensitive gFET arrays, enabled the development of an ultrasensitive and selective glutamate aptasensor. Upon in vitro validation using glutamate in artificial cerebrospinal fluid (aCSF), we demonstrate the aptasensor’s potential and usefulness in a clinical context by measuring glutamate concentration in CSF of AD patients. We also show, for the first time, that CSF glutamate concentration correlates with neurofilaments, an emerging biomarker of neuronal cell death.

## 2. Results and Discussion

### 2.1 In silico design and characterization of a novel glutamate-specific DNA aptamer

To obtain selective and ultrasensitive biosensors based on aptamers and FETs, the availability of short aptamer sequences with mobile sections that can reorient charges within the electrolyte’s Debye length is critical. Thus, to select a novel glutamate-specific aptamer that would be suitable as a molecular probe in gFET biosensors, we started by designing and characterizing in silico a shorter and more compact version of a previously described, 98 nt-long DNA aptamer termed glu1d04, with sequence 5’-GCATCAGTCCACTCGTGAGGTCGACTGATGAGGCTCGATCAGGAGCGCCGCTCGATCGCACTTTCACAGGATAGTAGTTGGTAGCGACCTCTGCTAGA-3’ [48]. In a first step, the secondary structure of this aptamer was predicted in silico using MFold [49] at 37 ⁰C and 25 ⁰C, the temperatures at which in vivo and in vitro experiments are usually performed, respectively (Suppl. Fig. 1A). The ionic conditions chosen were 100 mM Na^+^ and 1 mM Mg^2+^, as they correspond to the buffers typically used in aptamer-based biosensing. The most stable secondary structures (i.e., those showing the Minimum Free Energy, MFE) of glu1d04 show three to four elements that were formed and connected by open regions, with hairpins spanning nucleotides 2-30 and 36-58, at both temperatures (Suppl. Fig. 1A).

Based on these data, several strategies were tested to trim the glu1d04 aptamer to the shortest variant that could retain the two main hairpin motifs present in the original molecule at both 37 ⁰C and at 25 ⁰C, as they could be involved in the glutamate-binding site(s). Also, different point mutations, insertions or deletions that could increase the stability of the overall secondary structure were tested (data not shown). The best result was obtained by trimming 40 nt of the 3’ end of glu1d04, without introducing any additional nucleotide changes. The sequence of the resulting 58 nt-long DNA molecule, which we have named NG-Apt-Glu, is 5’-GCATCAGTCCACTCGTGAGGTCGACTGATGAGGCTCGATCAGGAGCGCCGCTCGATCG-3’. The secondary structure prediction of the aptamer (Fig. 1A), obtained with MFold, shows 18 unpaired nucleotides at different positions, including two hairpin loops, named site A (nucleotides 14 to 18) and site B (nucleotides 29 to 39). The prediction also describes an unconventional pairing of nucleotides between G13 and T19. Additionally, QGRS Mapper [50] was used to detect the potential formation of G-quadruplexes in the aptamer, but no such four-stranded DNA structural motifs were detected.

**Figure 1.**
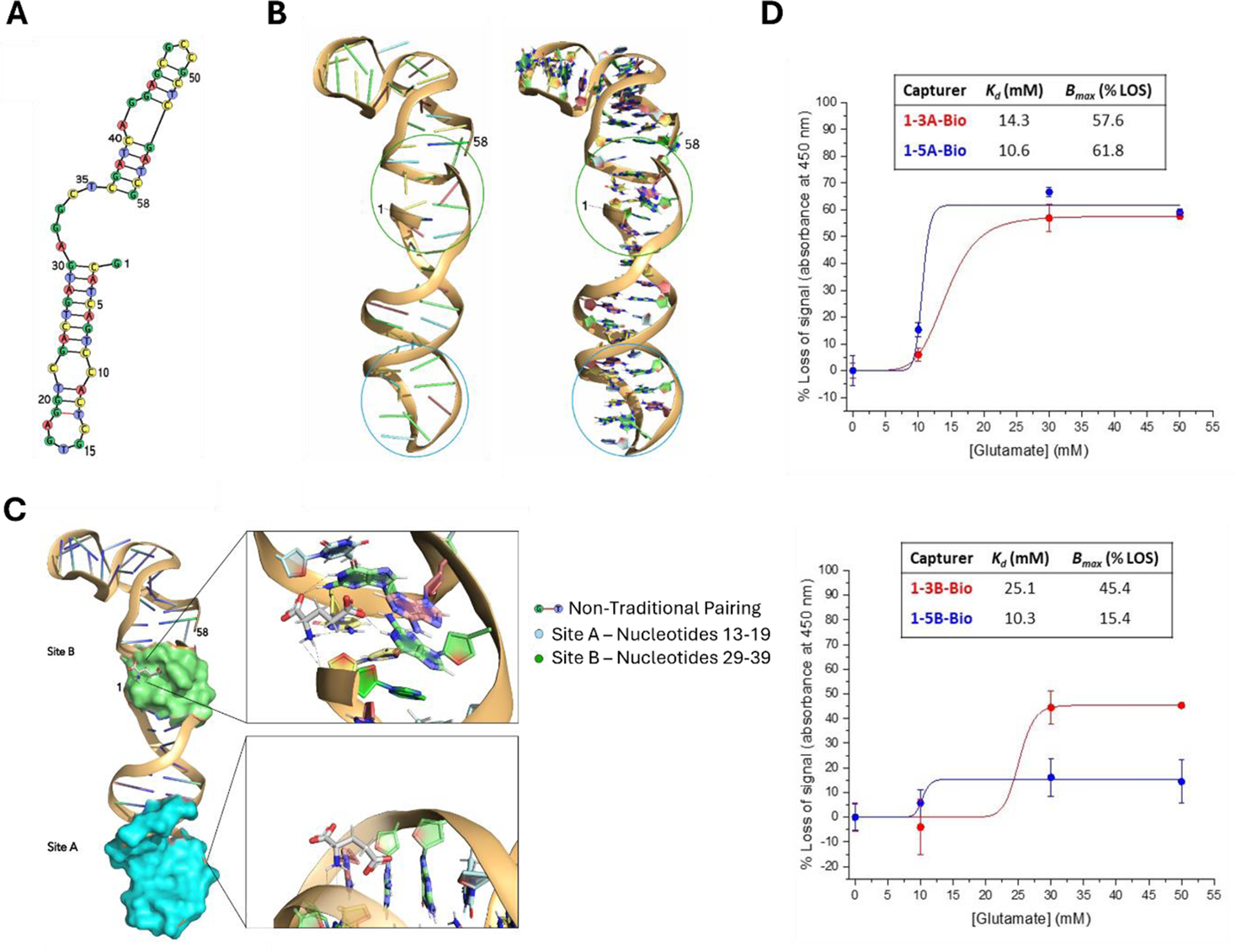
*In silico* design and simulations of the novel glutamate-specific DNA aptamer NG-Apt-Glu, and in vitro testing of its binding potential. **A)** Predicted secondary structure of the aptamer NG-Apt-Glu; **B)** Predicted 3D structure of the aptamer and its predominant conformation, with post MD refinement (left) and post MD refinement with highlighted nucleotides (right); **C)** In silico prediction of glutamate binding sites A (cyan) and B (lime), and predicted conformation of glutamate docked at site A (bottom insert) and B (top insert). **D)** Characterization of the binding affinities of the aptamer NG-Apt-Glu for glutamate, using colorimetric ELONAs (with the four designed capturers: 1-3A-Bio or 1-5A-Bio (top); and 1-3B-Bio or 1-5B-Bio (bottom). Each data point shows the mean ± SD (n = 3). The best fit curves corresponded to Hill function binding model (r^2^ > 0.96) in all cases, which were used to quantify the *K_d_* (mM) and % LOS or *B_max_* values shown in the inserts.

The putative 3D structure of NG-Apt-Glu was then predicted in silico with Bio.tools’ 3dRNA/DNA web server [51]. This showed that the segments containing paired nucleotides display helix-like structures, while unpaired regions promote exposure of nucleotides and a higher availability for potential interaction with glutamate at the two alternative binding sites A and B (Fig. 1B). To further refine the 3D prediction and obtain a more detailed perspective of the range of conformations that the aptamer can adopt in solution, molecular dynamics (MD) simulations were performed, followed by a cluster analysis of the most dominant conformations. In 60% of the trajectory, the aptamer exhibits a more stable screw-like shape, with its 5’ and 3’ ends in closer proximity (Fig. 1B). Despite the resulting structure displaying a tighter conformation, both putative glutamate binding sites A and B remain exposed with organized, unpaired nucleotides, for potential interaction with glutamate.

To confirm that glutamate could interact with any of the identified potential binding sites, an in silico aptamer-ligand docking analysis was performed to compare the ability of glutamate to bind to each site. Glutamate bound with comparable affinity to both sites, with the scoring function suggesting a slight stronger affinity to site B (33.86) with respect to site A (30.15), consistent with the predicted poses (Fig. 1C). The docking predictions for site B demonstrate that glutamate can nest within that aptamer cavity and that this interaction is mediated at a significant extent by the acidic side chain of glutamate (in particular, one of the oxygen atoms of the carboxyl group), which is stabilized by hydrogen bonding within the site, namely with G30 and A31. The amine group of glutamate also forms hydrogen bonds with site B, namely with C34 and with the backbone of C2. In turn, site A can also be recognized by glutamate, although with a slightly weaker binding than in site B. In this case, the aptamer-ligand interaction is mediated primarily by the lateral group of glutamate, while the positively charged amine group interacts both with the negatively charged backbone of G15 and form hydrogen bonds with A18 (Fig. 1C). The carboxyl group is mainly exposed, with little indication of interaction with aptamer components.

### 2.2 In vitro characterization of aptamer-glutamate binding

To determine which of the two identified binding sites in the new aptamer would bind more strongly to glutamate in a physiological scenario, in vitro colorimetric binding assays were performed. Based on the two predicted binding sites for our NG-Apt-Glu, four different capturer oligonucleotides were designed that contained complementary sequences either to those at the terminal loop spanning nucleotides 14-18 of the aptamer (site A) and its flanking regions (capturers named as 1-3A-bio and 1-5A-bio), or to those containing the open region at nucleotides 31-35 (site B) and its flanking regions (capturers 1-3B-bio and 1-5B-bio). In both cases, the numerals 3 or 5 indicate the biotinylation at their 5’ or 3’ ends (Suppl. Table 1 and Suppl. Fig. 2). Inhibition colorimetric enzyme-linked oligonucleotide assays (ELONAs) were conducted to calculate the dissociation constant (K_d_) and maximum loss of signal (% LOS or B_max_) of the aptamer NG-Apt-Glu for glutamate, using the four designed capturers (Suppl. Fig. 2). The % LOS is directly proportional to the concentration of glutamate, as expected in this assay (Suppl. Fig 3). In our experimental conditions, K_d_ values were in the millimolar range (from 10.3 to 25.1 mM) while % LOS ranged from 15.4 to 61.8 (Fig 1D). Overall, the results show that, upon the interaction with glutamate, NG-Apt-Glu desorbs slightly more readily (i.e., with lower K_d_ and higher % LOS values) when it was hybridized to the capturers 1-5A-Bio and (to a lesser extent) 1-3A-Bio (Fig 1D). This suggests that glutamate can bind to both sites of the aptamer, as shown by the in silico simulations (Fig 1C), though with a relative preference for site A. However, the collaboration of other unpaired regions or the implication of coaxial stacking of the helical elements of NG-Apt-Glu in the glutamate binding site cannot be entirely discarded.

### 2.3 Aptasensor Development and NG-Apt-Glu Validation

In previous works, gFETs have been shown to work as ultrasensitive transducers for biosensors [52]. Thus, to achieve ultrasensitive glutamate detection, an aptasensor was developed by combining the new NG-Apt-Glu aptamer as the biorecognition element with gFET arrays as the transducing platform.

#### 2.3.1 gFETs Fabrication

Our gFET arrays were fabricated on a 200 mm silicon wafer, building on our previously published methods for reproducible wafer-scale gFET fabrication [47]. Wafers contained 729 identically sized 5 × 5 mm^2^ gFET array chips (Fig. 2A). Each chip consisted of a transistor matrix with 32 independent drains, two different sources (each connected to 16 channels with independent drains), and two common co-planar gates (Fig. 2A). The graphene channel, connecting the source and the drain of each transistor, was 68 × 20 µm^2^ (W × L). For electronics interfacing, transistor array chips were wire-bonded to a custom printed circuit board (PCB) with two separate connectors that allow addressing groups of 16 transistors at the same time with a custom electronics platform (Fig. 2B).

**Figure 2.**
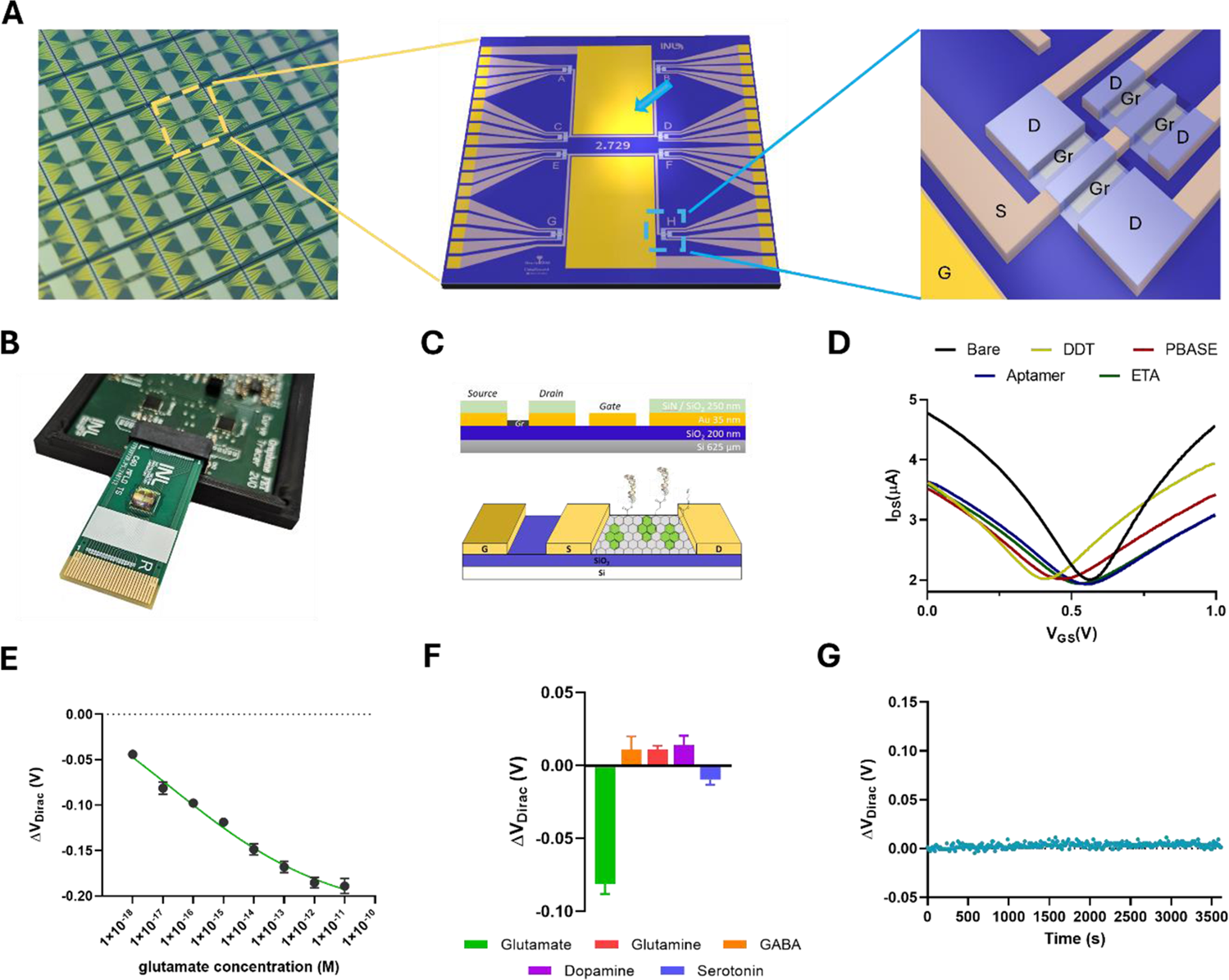
Aptasensor development and glutamate detection in artificial cerebrospinal fluid. **A)** Photograph of several diced gFET chips on the wafer (left); schematic illustration of the chip’s layout with mirrored sources and gate contacts on each side of the chip, featuring 32 drains in groups of four and two large co-planar gates (a blue arrow points to one gate) (center); and detail of a group of 4 gFETs with a single source (S) connected to 4 independent drains (D) by graphene channels (Gr) and partial view of the gate (G) (right); **B)** Photograph of a chip assembled on a PCB inserted in a custom-built electronics interface for data acquisition; **C)** Schematic illustrations of the gFETs fabrication layers with respective thicknesses (top; n.b. not to scale) and of a functionalized gFET with the glutamate-specific aptamer, linked to the graphene channel through a PBASE linker (bottom) (G: gate; S: source; D: drain; Au: gold; SiO_2_: silicon dioxide; SiN: silicon nitride; Si: silicon); **D)** Representative transfer curves from one gFET as measured in 1× aCSF after each biofunctionalization step; **E)** Calibration curve for glutamate detection in 1x aCSF for concentrations spanning from 1 aM (1×10^-18^ M) to 10 pM (1×10^-10^ M). **F)** Selectivity analysis of the glutamate aptasensor against molecules involved in the glutamate synthesis pathway (glutamine and GABA) and other neurotransmitters (dopamine and serotonin). Glutamate concentration is 10 aM and GABA, glutamine, dopamine, and serotonin are 1 pM. **G)** Stability analysis of the aptasensor functionalized with NG-Apt-Glu glutamate aptamer with aCSF continuously incubated for 1 hour and measurements taken every 10 seconds.

#### 2.3.2 gFETs Biofunctionalization with NG-Apt-Glu

The gFETs were converted into aptasensors through a functionalization process that allowed immobilization of our new glutamate aptamer on the graphene channel of each transistor and passivation of the gold gates (Fig. 2C). The latter was achieved with a self-assembled monolayer of dodecanethiol (DDT), previously shown to block adsorption of biomolecules in solution to the gold layer and being compatible with biosensing platforms [53], [54]. To immobilize aptamers on the graphene channel surface, the crosslinker 1-Pyrenebutyric acid N-hydroxysuccinimide (PBASE) was non-covalently bound to graphene through π–π stacking, preserving graphene’s crystalline structure [55]. Then, the 5’ amine terminal group added to the NG-Apt-Glu aptamer during synthesis reacted with PBASE’s free ester group, forming an amide bond and thus covalently conjugating the linker with the aptamer. PBASE linkers that did not bind to any aminated aptamer probe were blocked by ethanolamine (ETA). We have previously shown that this functionalization approach can successfully immobilize DNA probes, including aptamers, on graphene and prevent or reduce biofouling for biosensing applications [47], [56]. At each functionalization step, gFETs’ characteristic ambipolar transfer curve was acquired in phosphate-buffered saline solution (PBS) (Fig. 2D). Immobilization of the aptamer on the PBASE linker was confirmed by a characteristic positive shift of the charge neutrality point (V_DIRAC_) of the gFETs of approximately 100 mV. This results from the negatively charged phosphate backbone of DNA creating an image charge of opposite signal, which positively dope graphene, and is in line with prior observations of DNA immobilization on graphene [47], [57].

#### 2.3.3 Glutamate Detection in artificial cerebrospinal fluid

To determine concentration calibration curves for glutamate with our aptasensor in a biologically realistic buffer, measurements were performed in artificial cerebrospinal fluid (aCSF), which mimics the electrolytic profile and osmolarity of biological cerebrospinal fluid (CSF). Glutamate was prepared from a stock solution and diluted in 1x aCSF for final concentrations between 1 attomolar (1 aM, 1×10^-18^ M) and 10 picomolar (10 pM, 1×10^-11^ M). The addition of 1 aM of glutamate in aCSF to the aptasensor led to an average shift of -44 ± 2 mV, with a subsequent linearized response to increasing concentrations of up to 1 pM (1×10^-12^ M), showing a 24 mV/decade peak sensitivity (Fig. 2E). The observed linear working range of 160 mV spanned 6 orders of magnitude in concentration, only saturating at 1 pM. The shift of the transistor’s transconductance curve toward negative values is indicative of an increase of positive charges in the graphene channel vicinity, within the Debye length. Considering that the aptamer is much larger than glutamate, as can be seen from the in silico binding studies taken for its development (Fig. 1C), it was hypothesized that glutamate binding may lead to an aptamer’s conformational change that pushes negative charges of the aptamer’s DNA phosphate backbone away from the graphene channel. This behavior has been previously reported for an aptamer-based FET sensor for serotonin [33].

Our aptasensor achieves a limit of detection (LOD) of 1 aM, which is several orders of magnitude lower than the reported LOD for enzymatic and non-enzymatic glutamate biosensors previously reported, regardless of the transduction approach, as summarized in Table 1. The linear working range of our sensor, spanning six decades of concentration, is larger than most previous approaches, which typically range from two to four decades. Of note, the biosensors that reportedly achieved the lowest LODs for glutamate also used aptamers as the biorecognition element [48], [58]. One report described the use of a peptide aptamer linked to a terminal ferrocene that drives the current response of an amperometric system upon glutamate binding for detection in the nM range [59]. Although polypeptide aptamers are increasingly used in biosensors due to the nature of the interactions they can form with some targets [58], their large size prevents their use in FET-based biosensors for ultrasensitive detection. The lowest previously reported LOD for a glutamate biosensor (1.3 fM, 1.3 x 10^-15^ M), used a 39 nt-long DNA aptamer termed glu1, which is a truncated version of the 98 nt-long glutamate aptamer glu1d04 [48] from which our 58 nt-long aptamer NG-Apt-Glu was derived in this work.

**Table 1.** Comparison of different biosensors for glutamate detection.

| Type | Configuration | Detection method | LOD | Linear range | Sensitivity | Selectivity (tested against) | Human Samples | REF |
| --- | --- | --- | --- | --- | --- | --- | --- | --- |
| Non-Enzymatic Transconductance | Aptamer/Graphene FET | DNA Aptamer | 1 aM | 1 aM to 1 pM | 24 mV/decade | GABA, Glutamine, DA, Serotonin | CSF | **This work** |
| Non-Enzymatic Electrochemical Amperometric ACV | HS- Aptamer-Fc/Au electrode | DNA Aptamer | 1.3 fM | 0.01 pM to 1 nM | NR | GABA, Aspartic Acid, Tyrosine, L-Lactate | Serum | [48] |
| Non-Enzymatic Amperometric DPV | Aptamer/NPs/Au | Peptide Aptamer | 0.1 nM | 0.1 nM to 1 mM | 0.1572 $\mu\text{A}/\text{M}$ | DA | - | [59] |
| Non-enzymatic CV | NiO/GCE | Nanocrystals | 272 $\mu\text{M}$ | 1 to 8 mM | 11 $\mu\text{A}/\text{mM}\cdot\text{cm}^{-2}$ | AA, UA | - | [22] |
| Non-Enzymatic CV | Au electrode/polymer | Polymer | 1.7 $\mu\text{M}$ | NR | $0.86 \pm 0.12^\circ/\mu\text{M}$ | GABA | - | [26] |
| Non-Enzymatic CV | GluBP/AuNP/SPCE | GluBP | 0.15 $\mu\text{M}$ | 0.1 to 0.8 $\mu\text{M}$ | NR | AA, Aspartate | - | [23] |
| Enzymatic Amperometric CV | GC/Pt-MWCNTs/PPy/GluOx | GluOx | 0.88 $\mu\text{M}$ | 10 to 100 $\mu\text{M}$ | 723.08 $\mu\text{A}\cdot\text{cm}^{-2}/\text{mM}$ | AA, UA, DA, Glutamine, | Serum | [60] |
| Non-Enzymatic I-V | ZnO-RuO<sub>2</sub> NPs/GCE | Glassy Carbon Electrode | 96 pM | 0.1 to 0.01 mM | 5.42 $\mu\text{A}/\mu\text{M}\cdot\text{cm}^{-2}$ | AA, UA, Choline, DI-Methione, Glutathione, Gly, Methionin, Thiourea | Serum | [61] |
| Non-Enzymatic CV | CuO-MWCNTs/SPCE Electrode | Nanocomposite | 17.5 $\mu\text{M}$ | 20 to 200 $\mu\text{M}$ | 8500 $\mu\text{A}/\text{mM}\cdot\text{cm}^{-2}$ | AA, UA Glutamine | Blood, Urine | [62] |
| Non-Enzymatic I-V | GCE/CO<sub>3</sub>O<sub>4</sub>NSs | Glassy Carbon Electrode | 10 μM | 0.1 nM to 0.1 M | 9.5x10<sup>-5</sup> μA/μM·cm<sup>-2</sup> | Glutamine, Aspartate, Cystine, Glycine, Thiourea, Glutathione, Bilirubin | Serum, Urine | [63] |
| Non-Enzymatic CV Amperometric | Chitosan-derived carbon foam electrode | Cu ions | 0.001 μM | 0.001 to 1000 μM | 1.9 x10<sup>4</sup> μA/mM·cm<sup>2</sup> | AA, UA, DA, Glutamine, | - | [20] |
| Non-Enzymatic CV DPV | rGO/VS<sub>2</sub> | Electrode | 0.056 μM | 0.004 to 0.3 mM | 0.797 μA/mM·cm<sup>2</sup> | AA, UA, DA, Gluc, Gly | - | [64] |
| Enzymatic Amperometric | GluOx/CNT/screen printed electrode | GluOx | 10 nM | 0.01 to 10 μM | 0.72 ± 0.05 μA/μM | NR | - | [15] |
| Enzymatic CV DPV | GLDH/rGO/NiF | GLDH | 0.1 μM | 5 to 300 μM | 4.8 μA/μM·cm<sup>2</sup> | AA, UA, DA, Gly, Gluc | - | [13] |
| Enzymatic FSCV | GluOx-Chitosan | GluOx | NR | 25 μM to 1 mM | 33 ± 1 pA/μM | NR | - | [65] |
| Enzymatic CV | GluOx-modified Pt | GluOx | NR | 50 to 200 μM | 0.243 mM | NR | - | [14] |
| Enzymatic Electrochemical CV | Pt/GluOx-modified SiC@C nanowire | GluOx | 4.16 μM | NR | 0.24 pA/μM | NR | - | [66] |
| Enzymatic Amperometric CV | Ptelectrode/Polymer | GluOx | 220 nM | - | 2.16 nA mm<sup>-2</sup>/μM | AA | - | [67] |
| Enzymatic Amperometric | GluOx/nano-CP/Pt | GluOx | 0.1 μM | 0.2 to 100 μM | NR | NR | - | [68] |
| Enzymatic Amperometric | GluOx/Chitosan/Pt electrode | GluOx | 2.5 μM | 20 to 352 μM | 86.8 ± 8.8 nA/μM·cm<sup>-2</sup> | NR | - | [69] |
| Enzymatic Amperometric | GluOx/Nafion/PPy/Pt | GluOx | 1 μM | NR | 2.46 ± 0.48 pA/μM | AA, DA | - | [70] |
| Enzymatic Amperometric | GluOx/PtBlk | GluOx | 2 μM | 0 to 250 μM | 80 ± 10 nA/μM·cm<sup>-2</sup> | AA, DA | - | [71] |
| Enzymatic | GluOx/Pt microelectrode | GluOx | 524 nM | NR | -16.7 ± 0.8 μA/M | AA | - | [72] |
| Enzymatic Amperometric | GluOx/microsensor | GluOx | 10 μM | 10 to 100 μM | 3.4 ± 0.94 pA/μM | AA | - | [73] |
| Enzymatic Amperometric | GluOx/microelectrode | GluOx | 1 μM | NR | 32 mA/M·cm<sup>-2</sup> | NR | - | [74] |

Abbreviations used in this table: AA – Ascorbic acid; ACV – Alternating current voltammetry; AuNP – Gold nanoparticle; CNT – Carbon nanotube; Co<sub>3</sub>O<sub>4</sub>NSs – Cobalt oxide nanosheets; CuO-MWCNTs – Copper oxide multi wall carbon nanotubes; CV – Current voltammetry; DA – Dopamine; DPV - Differential pulse voltammetry; FET – Field-effect transistor; FSCV – Fast-scan cyclic voltammetry; GABA – Gamma-aminobutyric acid; GCE – Glassy carbon electrode; GLDH – Glutamate dehydrogenase; Gluc – Glucose; GluOx – Glutamate oxidase; Gly – Glycine; Fc – Ferrocene; I-V – Current-Voltage; LOD – Limit-of-detection; NiF – Nickel fluoride; NiO – Nickel oxide; Pt – Platinum; NPs – Nanoparticles; PtBlk – Platinum black; PPy – Polypyrrole; rGO – Reduced graphene oxide; SiC@C – Silicon carbide commercial nanowire; SPCE – Screen-printed carbon electrode; UA – Uric acid; VS<sub>2</sub> – Vanadium disulfide; ZnO NPs – Zinc oxide nanoparticle; ZnO-RuO<sub>2</sub> – Zinc oxide-doped ruthenium oxide; NR – Not Reported

To confirm that the sensitivity of our aptasensor is driven by the new NG-Apt-Glu aptamer, the response of our gFETs to glutamate with the NG-Apt-Glu aptamer was compared with that of our gFETs with the glu1 aptamer. NG-Apt-Glu on our gFETs not only lowered the LOD (1 aM vs 10 aM), but also increased the linear response range (1 aM to 1 pM vs 10 aM to 0.1 pM) and the sensor’s sensitivity (24 mV/decade vs 13 mV/decade) when compared with glu1 on our gFETs (Suppl. Fig. 4). Thus, the availability of two glutamate binding sites in our aptamer (A and B, Fig. 1) instead of only one (site A in aptamer glu1) seems to contribute to this unprecedented sensitivity. Additionally, the large surface-to-volume ratio and exceptional electronic mobility of gFETs, along with low noise-transduction due to a fabrication protocol that preserves graphene properties, also increased the sensitivity of our glutamate biosensor with respect to that previously reported with glu1 as probe molecule [48]. This result reinforces that both biorecognition molecule and transduction platform have to be designed in conjunction to push the limits of biosensors’ sensitivity.

The selectivity of the aptasensor to glutamate was evaluated against chemically similar molecules involved in glutamate’s synthesis pathways, such as GABA (gamma-aminobutyric acid) and glutamine, as well as against other neurotransmitters, including dopamine and serotonin (Fig. 2F). The average magnitude of the sensor’s response (V_DIRAC_ shifts) to a glutamate concentration of 10 aM was -81 ± 7 mV. In contrast, the response to 1 pM solutions containing GABA, glutamine, dopamine, or serotonin was 11 ± 7 mV, 11 ± 2 mV, 14 ± 6 mV, and -10 ± 3 mV, respectively. Despite being five orders of magnitude more concentrated than glutamate, the other potentially co-founding or co-occurring molecules led to negligible responses when compared to it. Testing the glutamate biosensor’s response to other biologically co-occurring and chemically similar molecules is fundamental for translating into real-world biosensing applications. These selectivity tests demonstrate that both the NG-Apt-Glu aptamer and the aptasensor based on it are highly selective for glutamate.

Another relevant characteristic of a biosensor for translational applications is its operational stability, which is critically influenced by design choices, including fabrication materials and operation modes [75], [76]. GFETs have been shown to display drift of their transfer curves during continuous, prolonged operation [77]. To assess drift, our aptasensor was incubated with a 40 µL drop of 1x aCSF for 1 hour, and the transistors’ response was measured every 10 seconds (Fig. 2G). The calculated average of V_DIRAC_ shifts over time was as low as 3.2 ± 2.7 mV. This observed time-dependent response was significantly below the observed average of 44 ± 2 mV in the transistor response to a 1 aM glutamate concentration. It should be noted that the repeated excitation of the gFETs for the V_DIRAC_ extraction at every 10 seconds, with a total of 360 acquired transfer curves over 1 hour, will produce more drift than under normal operation conditions, where a series of 20 transfer curves is acquired in under 1 second after sample incubation. Thus, it is expected that the real intrinsic aptasensors’ response variance will be even lower.

### 2.4 Glutamate quantification in human CSF

Although previously proposed glutamate biosensors were tested in human samples such as blood, serum, and urine (Table 1) [60], [61], [62], [63], they have not been assayed in cerebral spinal fluid (CSF) or correlated with clinical features of relevant brain disorders. Thus, to assess the translational potential of our sensor, glutamate quantification was performed in CSF samples from patients referred to the neurology outpatient clinic of University Hospital Center of São João (Porto, Portugal) due to cognitive complaints. Participants (n=16) were followed for at least a year in which they underwent thorough clinical evaluation including objective neurological exam, magnetic resonance imaging (MRI), computerized tomography (CT), and lumbar puncture for CSF collection in the context of their disease investigation (Fig. 3A). Participants were then diagnosed with Alzheimer’s Disease (AD, n=10) or excluded from having any neurodegenerative condition (n=6), the latter considered controls for this study.

**Figure 3.**
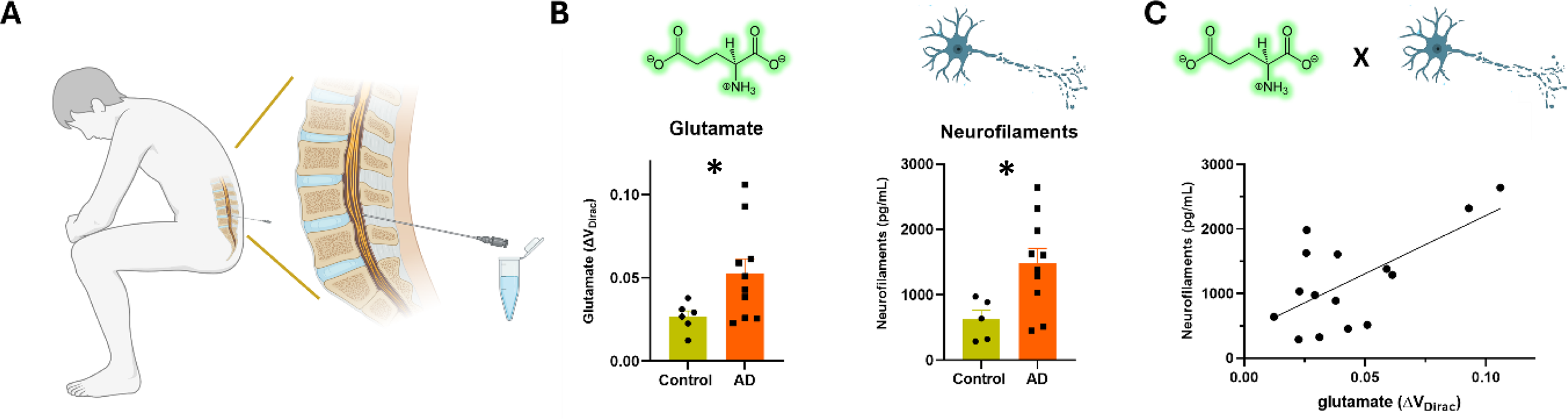
Glutamate and neurofilament light chain quantification in Alzheimer’s Disease fluid. **A)** CSF samples were collected from subjects referred to the neurology outpatient clinic for lumbar puncture in the context of their disease investigation, with a final diagnosis of Alzheimer’s Disease or no neurodegenerative disease (controls); **B)** Glutamate levels measured with our aptasensor (left) and neurofilament levels measured through ELISA (right) in CSF samples from Alzheimer’s Disease patients and controls (Kolmogorov-Smirnov, *p<0.05); **C)** Correlation between glutamate levels measured with the aptasensor and neurofilaments light chain (Nfl) measured with ELISA in the same patient samples (line is linear fit; Pearson correlation, r^2^=0.43, p=0.008).

Subsequently, 20 µL CSF samples from each participant were directly incubated in our sensors for 10 minutes, and transconductance was measured following rinsing. The quantified glutamate levels were on average two times higher in AD patients than in control subjects, with some AD patients showing glutamate levels 3-4 times higher than the controls’ average (Fig. 3B). Although the neurobiology of AD is not yet fully understood, there is accumulating evidence of disrupted glutamatergic neurotransmission [78], [79], [80], [81], and previous studies have also found increased glutamate levels in CSF of AD patients through HPLC [82], [83]. A critical difference between the measurements with our aptasensor and those reported with HPLC is that in our case no sample preparation processes were needed, and statistically significant results were obtained within 10 minutes.

A prevailing hypothesis for AD is that glutamate excitotoxicity leads to neuronal damage or cell death in multiple brain areas of the brain, including the hippocampus and frontal cortices, which correlates with cognitive and memory problems as well as neuroanatomical alterations found in AD patients [84], [85]. To assess a possible correlation between glutamate levels measured with our aptasensor and a marker for neuronal damage/death, neurofilament light chain (NfL) was quantified in the CSF samples from the studied patients. Neurofilaments are an integral part of the neuron’s cytoskeleton, and significant damage to the cells leads to an increase in neurofilament chains in CSF or plasma [86]. Hence, neurofilaments are an emerging biomarker for several neurodegenerative disorders [87], [88], [89], [90]. Measured NfL levels were on average two times higher in AD patients than in control subjects (Fig. 3B). In the studied patients, a significant correlation was found between the measured glutamate and NfL levels (r² = 0.43, p = 0.008) (Fig. 3C), underscoring the potential relationship between glutamate levels and cell death in AD. In the subset of control (n=5) and AD samples in which glutamate levels were above the control’s average (> 0.27, n=7), the significant correlation rises to r² = 0.73. Because different stages of AD, or even different onsets, may affect glutamate levels differently [85], further investigation of the correlation between glutamate and NfL levels in a larger segmented dataset is needed and will be analyzed in future studies. Nevertheless, as glutamate measurements were performed during clinical investigation and correlated with disease state, the quantification of this biomarker by means of our aptasensor can potentially contribute to the earlier diagnosis and staging of AD.

## 3. Conclusions

The graphene aptasensor for highly selective and sensitive glutamate detection proposed in this work combines an array of gFETs with a newly designed DNA aptamer. We show that combining in silico design and predictions with biochemical validation can accelerate the selection of optimal aptamers useful as molecular probes for biosensors. The conventional selection of novel aptamers via SELEX is a time-consuming process with a low success rate for small targets, such as neurotransmitters. In our approach, combining in silico and in vitro methods enabled deriving a new glutamate DNA aptamer, NG-Apt-Glu, with high affinity and specificity for glutamate.

The integration of the new aptamer into gFET arrays fabricated through reproducible clean-room methods led to the development of a glutamate aptasensor for detection and quantification of this essential neurotransmitter. The extraordinary sensitivity of our biosensing platform was driven by the extraordinary sensitivity of gFET sensors and was enhanced by the aptamer’s ability to drive transduction events at significantly lower glutamate concentrations and at a short distance from the graphene surface. The observed limit of detection of 1 aM of glutamate in aCSF, together with a wide linear working range from 1 aM to 10 pM, surpassed those of other recently published glutamate sensors, including those that use different aptamers or field-effect transistors. The potential contribution of two putative binding sites in our NG-Apt-Glu aptamer, identified in silico and validated in vitro, may also account for the increased sensitivity and wide working range observed. Our results show that both the biorecognition element and the transduction platform contribute to biosensor sensitivity and should be designed and optimized together in biosensor development.

The unprecedented sensitivity and specificity of our developed aptasensor in vitro suggested that it could detect small changes in the physiological concentration of glutamate. Thus, to assess its translational potential, glutamate was measured in the CSF of patients referred to Neurology consultations at a central hospital. Following diagnosis, it was found that the samples of patients diagnosed with Alzheimer’s Disease showed, on average, glutamate levels two times higher than those of control subjects. Additionally, we found that glutamate levels showed a significant correlation with neurofilaments light chain, an emerging biomarker of neuronal death in neurological disorders, in the same samples. This not only provides preliminary and significant observations on the potential correlation between the excitotoxic effects of excessive glutamate levels and neuronal survival, but also indicates that our aptasensors could be used as a promising tool for point-of-care diagnosis of neurological disorders. Future studies involving larger stratified patient pools, including those with neurological disorders other than Alzheimer’s, will be needed to validate the developed technology for clinical research and practice. This approach could also be extended to other neurotransmitters, using the appropriate aptamers, to further study the implications of imbalanced neurotransmitter levels across brain disorders that currently affect large fractions of the world population.

## 4. Materials and Methods

### 4.1. Materials and Reagents

Glutamate (L-glutamic acid, 99.9%), serotonin (5-hydroxytryptamine hydrochloride, 98%), dopamine (3,4-dihydroxyphenethylamine hydrochloride, 98%), GABA (gamma-aminobutyric acid, 99%), L-glutamine (99%), dimethyl sulfoxide (DMSO) (99.9% HPLC), 1-pyrenebutynic acid N-hydroxy-succinimide ester (PBASE, 95%), 1-dodecanethiol (DDT, 98%), ethanolamine (ETA, 98%), sodium chloride (NaCl), potassium chloride (KCl), monosodium phosphate (NaH_2_PO_4_), sodium bicarbonate (NaHCO_3_), glucose, calcium chloride dihydrate (CaCl_2_.2H_2_O), phosphate-buffered saline (PBS) tablets, poly(methyl(meth)acrylate) (PMMA) (5 k M.W. and 15 k M.W.), and sulfuric acid (H_2_SO_4_) were purchased from Merck. Anti-digoxigenin antibody labelled with horseradish peroxidase (HRP), bovine serum albumin (BSA), and Tris HCl (pH 7.6), were purchased from Sigma. Phosphate 10x buffer saline (PBS), tween 20, 3,3’,5,5’-tetramethylbenzidine (TMB), magnesium chloride (MgCl_2_), and calcium chloride (CaCl_2_) were purchased from ThermoFisher. Streptavidin coated multiwell plates were purchased from

Pierce. Nuclease-free water was purchased from BioLabs. Acetone (99.5%), ethanol (99.8%), and 2-propanol (99.8%) were purchased from Honeywell. Photoresist AZ1505 and AZ4110 were purchased from MicroChemicals GmbH. 25 µm thick high-purity (99.999% purity) copper (Cu) foils were purchased from Goodfellow. RTV silicone elastomer (3140, Dowsil) and superglue (Loctite) were acquired from Farnel. NG-Apt-Glu aptamer (5′-GCATCAGTCCACTCGTGAGGTCGACTGATGAGGCTCGATCAGGAGCGCCGCTCGATCG-3′), with a 5′ C6-amino modification, was synthesized by Stab Vida. Capture strands (1-3A-bio, 1-5A-bio, 1-3B-bio or 1-5B-bio), as well as the unlabeled aptamer NG-Apt-Glu, were synthesized by IDT.

### 4.2. In silico characterization of the 3D structure of the aptamer and its binding to glutamate

The pipeline for generating the 3D structure of the DNA aptamer followed a commonly used methodology [91], successfully employed in previous works [46], [92], [93]. Secondary structure was predicted with UnaFold web server [49]. To complement and further optimize the predictions, the temperature was set to 37 °C, while Na^+^ and Mg^2+^ concentrations were set to 100 mM and 1 mM, respectively. The threshold to limit folding was defined within 5% of the minimum free energy, which resulted in 50 foldings. There were no imposed limits for distances between paired bases.

The resulting secondary structure was retrieved and used as an input for predicting the aptamer 3D, tertiary structure using the 3dRNA/DNA tool [51]. This tool uses a template-based or a distance-geometry method to model 3D conformations from the secondary structure. The model outputs 10 structures scored through an internal scoring system. The best scoring structure was selected and downloaded in PDB format. With the goal of exploring the conformation flexibility of the aptamer under the effect of an explicit solvent, a molecular dynamics (MD) simulation was performed. Using the LEAP module of the AMBER20 [94] software package, the process starts with solvation of the aptamer structure using a TIP3P [95] rectangular box of water molecules. The box size guarantees that each atom is at least 12 Angstrom (Å) away from the edge. To achieve physiological concentration of NaCl within the simulation, the SPLIT [96] method was used, allowing calculation of the total number of Na^+^ and Cl^-^ ions, and targeting an ionic concentration of 0.15 M. The system was described with the BSC1 force field [97], developed for atomistic simulations of DNA. After solvation a sequence of minimization stages was performed, as to remove atomic clashing during the MD: water molecules for 2,500 steps, hydrogen atoms for 2,500 steps, side chains of the DNA structure for 2,500 steps and the complete system for 10,000 steps.

After concluding the minimization, the system was equilibrated through an MD procedure segmented in two steps. The first one begins with gradual heating of the system from 0 K up to 310.15 K (37 °C) for a total of 50 ps. Afterwards the system is equilibrated in terms of density for 50 ps while maintaining the temperature (310.15 K) constant. The final stage consists of a production run of 1000 ns using an NPT ensemble, with constant temperature and pressure at 310.15 K (Langevin thermostat) and 1 bar, respectively. With the goal of constraining covalent bonds involving hydrogen atoms, the SHAKE algorithm [98], [99], [100] was employed, together with a 2-fs time-step. The cutoff for the non-bonded interactions was set to 10 Å and kept throughout the simulation. The conformations and trajectories were analyzed through backbone Root Mean Square Deviation (RMSD) using CPPTRAJ [101], from which the most prevalent conformations were retrieved.

GOLD software [102] was employed to evaluate glutamate ability to bind to the NG-Apt-Glu aptamer, allowing a detailed exploration of the putative binding sites A (nucleotides 13 to 19: 5’-TCGTGAG-3’) and B (29 to 39: 5’-TGAGGCTCGAT-3’). GOLD applies a non-deterministic genetic algorithm (GA) to generate ligand poses and employs a previously selected scoring function to evaluate the affinity between the two molecules. The exploration space included the nucleotides defined for sites A and B, plus a 30 Å radius. The scoring function PLP score was applied with a total number of 100 GA runs. The resulting scores and most stable poses were compared for each site.

### 4.3. Biochemical characterization of the glutamate aptamer by inhibition colorimetric ELONAs

The dissociation constant (*K_d_*) and maximum loss of signal (% LOS or *B_max_*) of the aptamer NG-Apt-Glu were determined by inhibition colorimetric Enzyme-Linked Oligonucleotide Assays (ELONAs). Based on our previous experience with ELONA methodology [103], [104] and the features of this inhibition strategy [105], 1.25 pmoles of biotin-tagged capturer DNA oligonucleotides (shortened as “capturers” and named as 1-3A-bio, 1-5A-bio, 1-3B-bio or 1-5B-bio, Suppl. Table 1 and Suppl. Fig. 2), were resuspended in 100 µl of selection buffer (SB) (100 mM NaCl, 20 mM Tris HCl pH 7.6, 5 mM KCl, 2 mM MgCl_2_, 1 mM CaCl_2_) and immobilized onto pre-blocked (with bovine serum albumin, BSA) streptavidin-coated multiwell plates by incubating at 25 °C for 1 h, with agitation of 350 rpm. Non-adsorbed capturers were removed by washing the wells three times with 100 µl of SB. In parallel, 1.25 pmoles of the 5’ digoxigenin-labelled ssDNA aptamer NG-Apt-Glu (or “tracer” for the ELONA, Suppl. Table 1 and Suppl. Fig. 2) were resuspended in 100 µl of SB and subjected to a denaturation/renaturation process through three consecutive incubations at 90 °C, 4 °C and 25 °C, for 10 min each and with agitation of 350 rpm. The renatured aptamer was added to each well containing the corresponding biotin-labeled capturer (Suppl. Fig. 3). Then, the plate was incubated at 25 °C for 2 h with agitation of 350 rpm, and the aptamers that did not bind to the capturers were discarded by washing the wells two times with 100 µl of SB (the first one at 28 °C and the second one at 25 °C) for 15 min at 350 rpm.

The aptamer-capturer complexes that remained bound to the microplate wells were incubated with 100 µl of glutamate at four different concentrations (0, 10, 30 and 50 mM in SB) at 25 °C for 45 min, with agitation of 350 rpm. To quantify the digoxigenin-labelled aptamers that remained bound to the capturers onto the microplate surface, 100 µl of an anti-digoxigenin antibody conjugated to horseradish peroxidase (HRP) (diluted 1:5000 in 1x

PBS supplemented with 1% BSA, 1 mM MgCl_2_ and 0.1% Tween 20) were added to each well and incubated at 37 °C for 30 min at 350 rpm. After washing three times with 200 µl of SB, 100 µl/well of the chromogenic substrate 3,3’,5,5’-tetramethylbenzidine (TMB) were added and incubated at 25 °C for 5 min. The HRP-catalyzed reaction was stopped by adding 100 µl/well of 2 M H_2_SO_4_. The absorbance at 450 nm was measured in each well using a microplate reader Tecan GENios (Tecan Group Ltd.). The loss of signal (LOS) quantified in the wells where glutamate solutions were added, with respect to the signal measured in those without added glutamate (100 % of the signal, corresponding to 0 mM glutamate), showed that the desorption of the aptamers from the capturers was triggered by the presence of glutamate in the medium, being % LOS and [glutamate] directly proportional (Suppl. Fig. 3). The binding curves were fitted using a sigmoidal Hill function and the Levenberg-Marquardt iteration algorithm, using Origin 2021 (Mathworks). The dissociation constant (*K_d_*) and maximum % LOS or *B_max_* for each combination of capturer-aptamer were quantified by this software (Fig. 1D).

### 4.4. gFETs Fabrication

Chemical vapor deposition (CVD) was utilized to grow single-layer graphene on 25 µm-thick high-purity (99.999% purity) Cu foils in a three-zone quartz tube furnace (EasyTube ET3000, CVD Corp.). Cu substrates were treated with a mixture of FeCl_3_, HCl, and deionized water for 2 min in ultrasound and then placed in the furnace. A 300-sccm Ar and 120-sccm H_2_ gas mixture was pumped into the system after it had been evacuated to 2 mTorr. 1.25 sccm of CH_4_, the carbon precursor, was added to the chamber when the growth temperature and pressure were reached. The growth was conducted for 45 minutes at 1040 °C and 6 Torr. After growth, graphene grown on the top side of the copper was protected by a poly(methyl methacrylate) (PMMA) mixture, and oxygen plasma ashing was performed to remove the graphene from the backside of the substrate. Raman spectroscopy was used to assess the quality of the grown graphene, as previously reported by us [47], [57].

A 200 mm Si wafer doped with B and with 200 nm thermal SiO_2_ was used as the substrate for the wafer-level production of the gFETs. Au (35 nm), the conductive layer, was sputter-coated onto the wafer with Cr (3 nm) as the adhesive layer, and with Al_2_O_3_ (20 nm) on top as a protective layer to avoid the gold alloying in further steps. Optical lithography and ion milling were used to pattern the source, drain, and gate Au electrodes. As a protective layer for the regions where the transferred graphene would be etched, a sacrificial layer (TiWN, 5 nm; AlSiCu, 100 nm; TiWN, 15 nm) was sputtered and patterned by lift-off. The previously grown graphene was transferred to the wafer through a PMMA-assisted wet transfer method [106] and the graphene channels were patterned via optical lithography and etched by reactive ion etching with oxygen plasma. Wet etching was then used to remove the sacrificial layer. A protective layer of Ni (100 nm) was sputtered and patterned by lift-off on the graphene and gate areas to function as a stopping layer for the passivation layer etch by ion milling. A multistack passivation layer of alternated SiO_2_ (50 nm) and SiN_x_ (50 nm) (total thickness of 250 nm) was grown via plasma-enhanced CVD, patterned via optical lithography and etched to cover the entire sensor area except for the graphene channel and contact gate area. Finally, wet etching of the protective layers, Ni and Al_2_O_3_, was performed. The wafer was diced into 729 equal-sized 5 × 5 mm^2^ chips after spin-coating a protective layer of photoresist which was later dissolved in acetone and then further rinsed with IPA and DI water after the dicing. The chips were glued onto custom printed circuit boards (PCB) with superglue (Loctite) and bonded with 25 µm-thick gold wires (wire bonder HB16 TPT HB) which were then protected with a silicon elastomer (3140, Dowsil).

### 4.5. gFETs Biofunctionalization

The gold gate on the gFET chips was first passivated with the incubation of 20 µL of 2 mM 1-Dodecanethiol (DDT, 2 mM in ethanol) in a humid chamber for 2 hours, followed by rinsing with deionized water and drying under N_2_ flow. Then, gFETs were incubated for 2 hours in the dark with 20 µL of the crosslinker 1-pyrenebutyric acid N-hydroxysuccinimide (PBASE, 10 mM in dimethyl sulfoxide, DMSO). The aptamer NG-Apt-Glu DNA, synthesized with a 5′ amino-link termination, was diluted to 20 µM in nuclease-free water. The solution was heated to 95 °C for 20 minutes and cooled down to room temperature. Then, 20 µL aptamer solution was incubated on the gFETs for 16 hours. Any PBASE on graphene’s surface that did not bind to the aptamers was blocked by incubation with 20 μL of ethanolamine (ETA, 100 mM in DI water) for 30 min. Finally, the gFETs were dried under N_2_ flow and rinsed with deionized water. At each step of the functionalization process the transconductance of each gFET was measured by applying a source-drain voltage (V_DS_) of 15 mV and measuring source-drain current (I_DS_) in gate-source voltage (V_GS_) sweeps between 0 and 1V.

### 4.6. Glutamate detection in aCSF

Glutamate in powder form was prepared into a stock solution of 10 mM in artificial cerebrospinal fluid (aCSF) (119 mM NaCl, 2.5 mM KCl, 1.2 mM NaH_2_PO_4_, 24 mM NaHCO_3_, 12.5 mM glucose, 2 mM MgSO_4_.7H_2_O, and 2 mM CaCl_2_.2H_2_O, with 300–310 mOsm/L) at pH 7.2–7.4. The stock solution was then diluted in 1 x aCSF to prepare solutions with varying glutamate concentrations from 1 aM (1×10^‒18^ M) to 10 pM (1×10^‒11^ M). For transconductance measurements, a custom-designed signal acquisition platform that can read 16 transistors sequentially was connected to the aptasensor chips via the custom PCB. gFETs baseline transconductance was measured in glutamate-free 1x aCSF by applying a V_DS_ of 15 mV and sweeping the V_GS_ between -0.4 and 0.6 V. Following baseline measurements, 20 μL samples of each glutamate concentration were incubated in the aptasensor for 1 h. After incubation of each concentration, gFETs were rinsed with aCSF and transconductance measured in 20 µL of 1 x aCSF with the same voltage parameters used for baseline measurements. ΔV_DIRAC_ was calculated as the voltage difference at which the Dirac point was observed between the baseline and each incubated glutamate concentration. For the glutamate concentration calibration curve, ΔV_DIRAC_ for each concentration was averaged from ten aptasensor chips measurements.

For the specificity experiments, GABA, glutamine, serotonin, and dopamine in powder form were diluted in 1 × aCSF to obtain 1 pM solutions. 20 μL samples of each solution were incubated in the aptasensors for 1 h. Following rinsing with aCSF, transconductance measurements were performed in 1 × aCSF using the previously mentioned voltage parameters. Transistor measurements from at least four aptasensors contributed to the average ΔV_DIRAC_ values reported for each solution.

### 4.7 Glutamate detection in patient CSF samples

Subjects were referred to the Neurology Service of the University Hospital Center of São João (CHUSJ, Porto, Portugal) due to cognitive complaints. All subjects underwent an evaluation comprising medical history, physical and neurological examination, neuropsychological assessment, CSF and blood collection, and brain CT and/or MRI scan. CSF was obtained from each participant by a lumbar puncture performed in the lateral decubitus position, at L4/L5 or L3/L4 interspace, under sterile conditions. CSF was collected in sterile polypropylene tubes, and aliquots of 500µl were transferred to Eppendorf tubes, which were stored at -80°C until further use. Following clinical investigation, subjects were either diagnosed with Alzheimer’s Disease (n=10) or excluded from having a confirmed neurodegenerative condition. The latter were considered as control subjects (n= 6) for the purpose of this study.

For glutamate quantification in CSF with our aptasensors, baseline transconductance was initially measured in 1x aCSF with the same voltage excursion parameters described above. Then, each CSF aliquot was thawed in ice and 20 µL samples were directly incubated in the chips for 10 min. Following rinsing with aCSF, transconductance was measured in 20 µL 1x aCSF with the same voltage parameters used for baseline acquisition. ΔV_DIRAC_ between baseline and sample transconductance curves were calculated for each sample.

In parallel, neurofilaments light chain (Nfl) were quantified in the same CSF samples using the NF-light^TM^ ELISA assay kit (UmanDiagnostics AB) according to manufacturer instructions. In one of the control samples, it was not possible to quantify Nfl concentration.

### 4.8. Statistical analysis

Statistical analysis was performed with GraphPad Prism 10 for Windows (Graphpad Software). Statistical details can be found in figure legends. Data in graphs are mean ± SEM, except where noted.

## Acknowledgements

The authors would like to thank: Ana Carolina Castro and Margarida Falcão of the Neurophysiology and Neuroengineering Lab (FMUP/Rise-Health) and Paula Serrão of the Pharmacology and Therapeutics unit (FMUP/ Rise-Health) for help in the preliminary test measurements with human samples; María Fernández-Algar (CAB) for her technical help with in vitro validation of the aptamer; and Tiago Pereira of the Alpuim group (INL) for the transistor schematics image. This work was funded by “la Caixa” Banking Foundation under the grant agreement LCF/PR/HR21-00410 (“NeuralGRAB”), and COMPETE2030-FEDER project 2023.17917.ICDT. Work at CAB was also supported by grant No. PID2022-139908OB-I00, funded by the Spanish Ministry of Science and Innovation / State Agency of Research MCIN/AEI/FEDER 10.13039/501100011033 and by “ERDF A way of making Europe”. Work at BioSim received financial support from portuguese national funds (FCT/MECI, Fundação para a Ciência e Tecnologia and Ministério da Educação, Ciência e Inovação) through project UID/50006 – “Laboratório Associado para a Química Verde - Tecnologias e Processos Limpos” – and Centro Nacional de Computação Avançada (CNCA) funded by FCT. MA was supported by FCT doctoral fellowship 2022.14536.BD. Some figures were created with Biorender.com.

## Author contributions

YB and CB designed the new aptamer in silico and developed the in vitro protocol to test its binding efficiency. RPG and SFS performed computational characterization of the aptamer. YB performed biochemical characterization of aptamer. MA and JB fabricated the sensors. MA performed functionalization and experiments in vitro and with human samples. IM and MVS coordinated patient sample and clinical data collection. IM performed neurofilaments quantification and analyzed patient data. MA and LJ analyzed aptasensor data. PM, JB, MVC, SFS, CB, LJ, and PA supervised the work and interpreted results. MA, YB, RPG, CB, and LJ wrote the first draft of the manuscript. SFS, IPM, MVC, CB, LJ, and PA revised the manuscript. All authors read and approved the final version of the manuscript.

## Ethics statement

Ethical approval was obtained from the ethics committee at University Hospital Center of São João (approval number 316/19). All participants provided their written informed consent prior to participating. The study conformed to the standards set by the Portuguese Law 21/2014 regulating clinical research, the EU’s General Data Protection Regulation (GDPR) 2016/679, and the Declaration of Helsinki.

