## Supplementary Material for "Ultrasensitive graphene FET aptasensor for direct attomolar detection of glutamate in human clinical samples"

**Suppl. Fig. 1: Secondary structure of glu1d04 and NG-Apt-Glu aptamers**

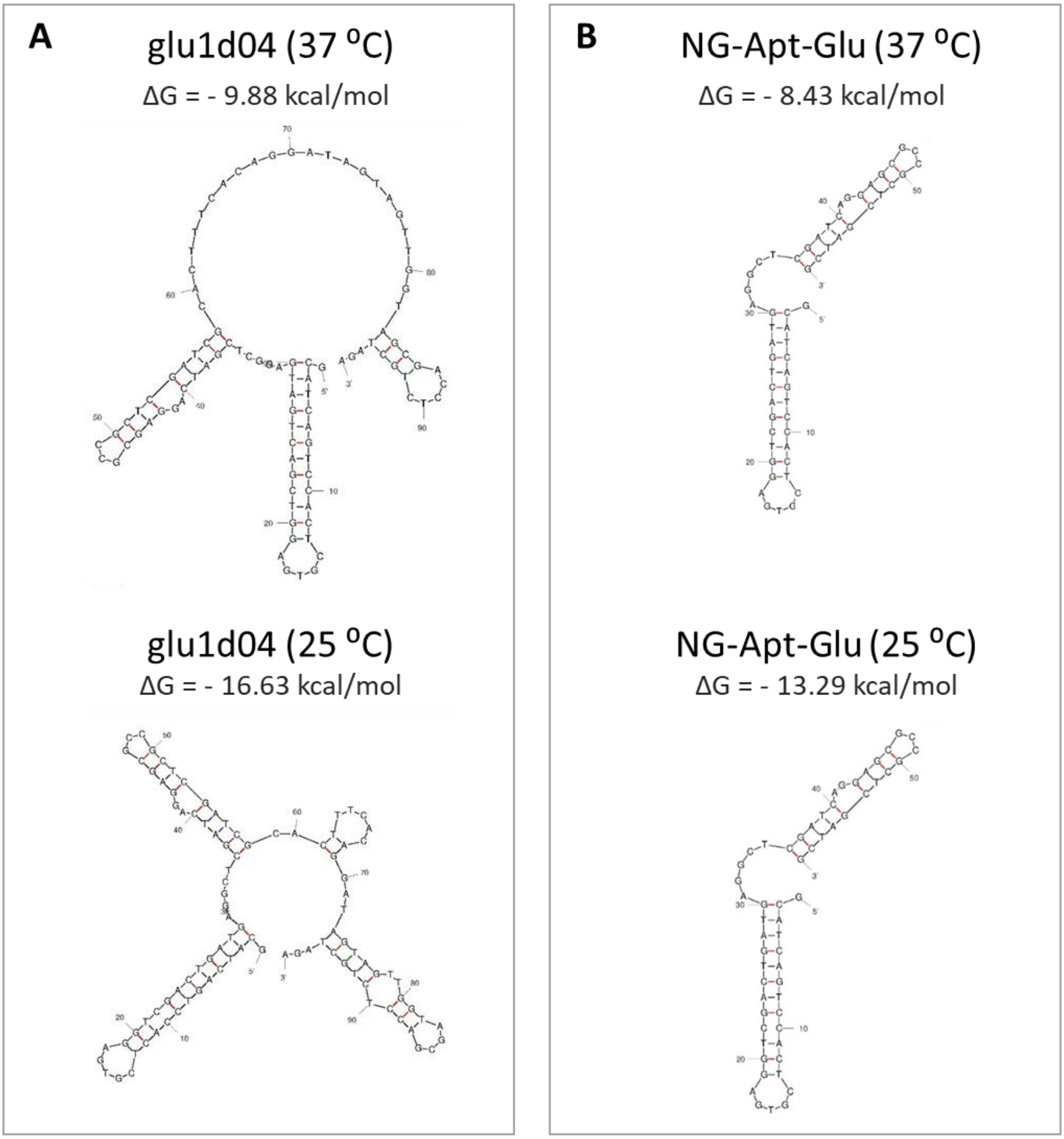


**Suppl. Fig. 1.** Secondary structures folded in silico using Mfold software [1] and their associated Minimum Free Energy (MFE, in kcal/mol), at 37 ºC and 25 ºC, of aptamer glu1d04 (described by Wu et al. [2]) (**A**), and the newly designed aptamer NG-Apt-Glu (**B**).

**Suppl. Fig. 2: Sequence and binding positions of four different capturer oligonucleotides for the two predicted binding sites of glutamate in aptamer NG-Apt-Glu**

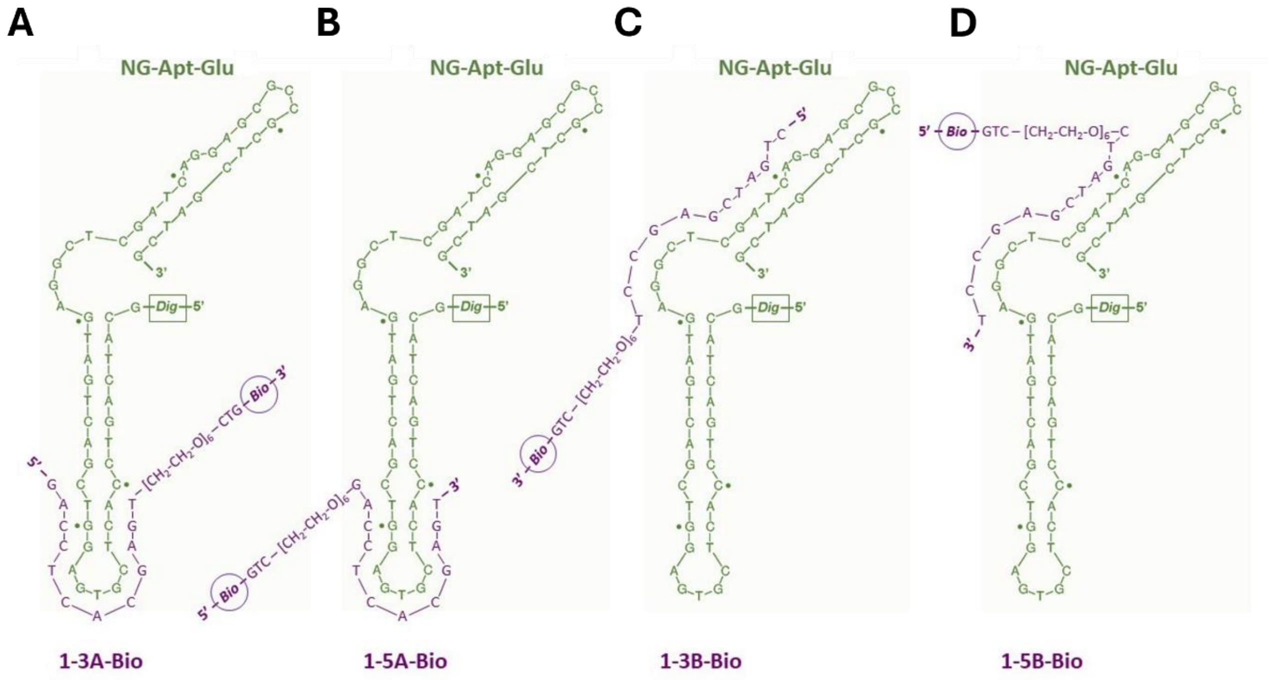


**Suppl. Fig. 2.** Secondary structures of the aptamer NG-Apt-Glu (in green) and the four capturers (in purple) used in inhibition colorimetric ELONAs: **A**) 1-3A-Bio; **B**) 1-5A-Bio; **C**) 1-3B-Bio; **D**) 1-5B-Bio. The expected hybridization positions through their complementary sequences (underlined in Suppl. Table 1) are shown.

**Suppl. Fig. 3: Process of the inhibition colorimetric Enzyme-Linked Oligonucleotide Assay (ELONA)**

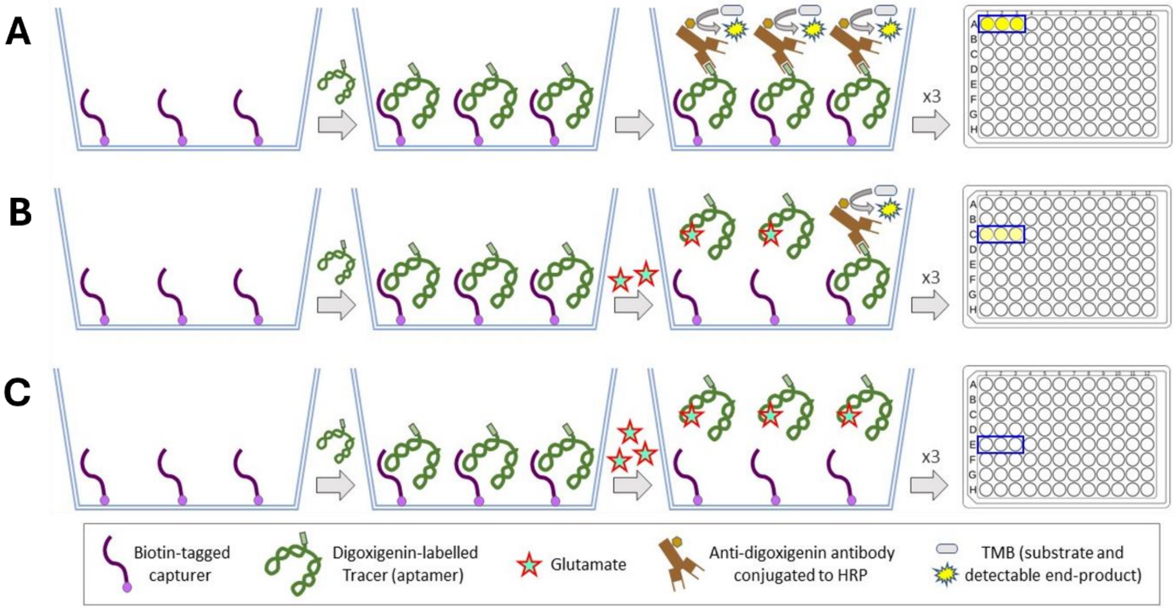


**Suppl. Fig. 3.** Scheme of the stepwise process of the inhibition colorimetric Enzyme-Linked Oligonucleotide Assay (ELONA) used in this study to measure the dissociation constant ($K_d$) and maximum loss of signal (% LOS or $B_{max}$) of the aptamer NG-Apt-Glu: **A**) illustrates the process without added glutamate, resulting in 100% signal (or 0% LOS); **B**) and **C**) show the effects of increasing concentrations of glutamate.

**Suppl. Fig. 4: Glutamate detection with glu1 aptamer in gFET aptasensors**

**A** **B**

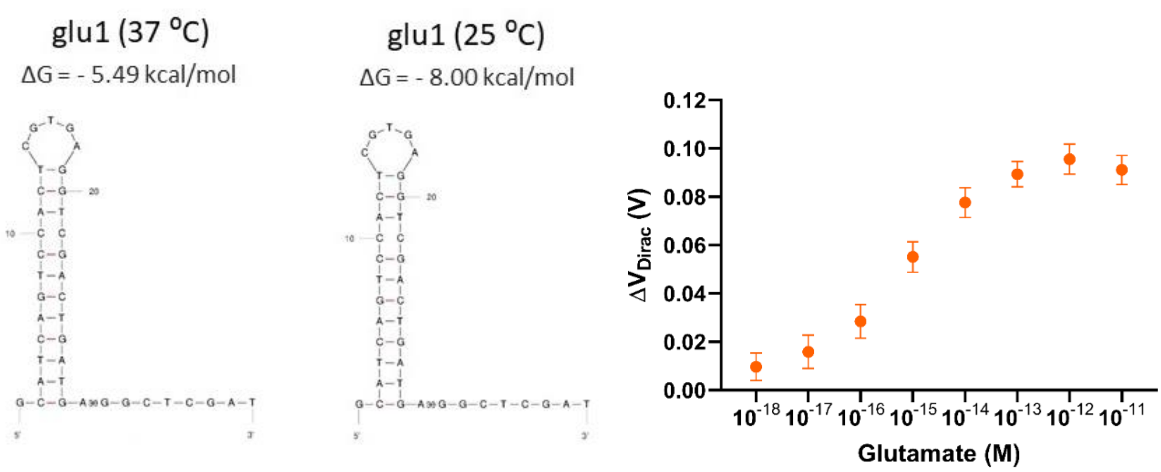


**Suppl. Fig. 4. A**) Secondary structures (folded in silico using Mfold software [1]) and their associated Minimum Free Energy (MFE, in kcal/mol), at 37 °C and 25 °C, of aptamer glu1 [2]. **B**) Calibration curve of glutamate detection in 1 x aCSF using our gFET arrays functionalized with aptamer glu1.

**Suppl. Table 1.** Sequences of the aptamer NG-Apt-Glu designed in this work and used as tracer in the inhibition colorimetric ELONAs performed, as well as those of the four capturer oligonucleotides designed to interact with it. Complementary, 12 nt-long sequences are underlined in tracer and capturers. Abbreviations: *Dig*, digoxigenin; *Bio*, biotin (which is bound to the spacer GTC-[$CH_2$-$CH_2$-O]$_6$).

| **Name** | **Nucleotide sequence (5'-3')** |
| --- | --- |
| NG-Apt-Glu | *Dig*-GCATCAGTCCACTCGTGAGGTCGACTGATGAGGCTCGATCAGGAGCGCCGCTCGATCG |
| 1-3A-Bio | GACCTCACGAGT-*[$CH_2$-$CH_2$-O]$_6$*-CTG-*Bio* |
| 1-5A-Bio | *Bio*-GTC-*[$CH_2$-$CH_2$-O]$_6$*-GACCTCACGAGT |
| 1-3B-Bio | CTGATCGAGCCT-*[$CH_2$-$CH_2$-O]$_6$*-CTG-*Bio* |
| 1-5B-Bio | *Bio*-GTC-*[$CH_2$-$CH_2$-O]$_6$*-CTGATCGAGCCT |
